# Noradrenergic modulation of stress induced catecholamine release: Opposing influence of FG7142 and yohimbine

**DOI:** 10.1101/2024.05.09.593389

**Authors:** Vladimir Visocky, Carleigh J. Turner, Matthew H Lowrie, Anthony Alibro, Fany Messanvi, Yogita Chudasama

## Abstract

**Background**: Life stress modulates decision making, particularly in the face of risk, in some cases prompting vulnerable populations to make suboptimal, life-altering choices. In the brain, stress is known to alter the extracellular release of catecholamines in structures such as basolateral amygdala (BLA) and nucleus accumbens (NAc), but the relationship between catecholamines and decision-making behavior under stress has not been systemically explored.

**Methods:** We developed an operant touchscreen decision-making task for rats comprising elements of loss aversion and risk seeking behavior. Rats were first injected systemically with an adrenergic α2A-receptor agonist (guanfacine) and antagonist (yohimbine), as well as a partial inverse GABAA agonist, FG 7142, known to induce anxiety and stress related physiological responses in a variety of species, including humans. We then used fiber photometry to monitor NE in the basolateral amygdala (BLA), and DA activity in the nucleus accumbens (NAc) while animals engaged in decision-making and following systemic injections of FG 7142 and yohimbine.

**Results:** Neither yohimbine nor guanfacine had any impact on decision making strategy but altered motivational state with yohimbine making the animal almost insensitive to the reward outcome. The pharmacological induction of stress with FG 7142 biased the rats’ decisions towards safety, but this bias shifted toward risk when co-treated with yohimbine. In the BLA and NAc, the FG 7142 altered catecholamine release, with systemic yohimbine producing opposing effects on NE and DA release.

**Conclusions**: Stress induced changes in catecholamine release in the BLA and NAc can directly influence loss sensitivity, decisions and motivation, which can be modulated by the α2A adrenoreceptor antagonist, yohimbine.

## INTRODUCTION

Everyday decisions involve inherent uncertainty or insufficient knowledge to make informed choice [1]. Such decisions often allow for choices that vary in their ‘risk,’ with one option being relatively safe and another that offer an uncertain potential for major gains and losses. In animals, as in humans, the decision-making is naturally biased towards risk aversion, a default behavioral mode observed in species as diverse as fish [2, 3], birds [4, 5] and bumblebees [6, 7]. Risk aversion is thought to be driven by the affective consequence of loss, also called ‘loss aversion,’ where a given amount of loss impacts humans and other animals more severely than experiencing an equivalent gain [8]. In humans, loss aversion is sensitive to stress [9–11], and patients with neurological or psychiatric illnesses are particularly vulnerable to the detrimental effects of stress thought to cause suboptimal or life-altering choices with long-term negative consequences [12–14]. Here we investigate the effect of stress-induced neuromodulation on risky decision making and brain regions that modulate loss aversion in rats.

Many acute stressors increase extracellular concentrations of norepinephrine (NE) and dopamine (DA) in a number of brain regions, indicating that stress can elicit widespread activation of catecholaminergic neurons [15]. Stress-susceptible brain regions that exhibit structural and/or functional alterations include the amygdala and nucleus accumbens [16–18], both of which contribute significantly to decisions that involve aligning reward gain with loss-sensitivity [19–23]. Stressful stimuli increase NE levels in the basolateral amygdala [24, 25] which produces inhibitory effects through α2 receptors to promote the stress response [16, 26, 27]. Thus, NE- modulation of stress in the amygdala could have a role in modulating loss aversion during decision making through stimulation of α2 receptors [19, 28]. Midbrain neurons containing and secreting dopamine (DA) are also activated during stress [29–31] causing a change in motivational state mediated by DA in the nucleus accumbens [32]. Consequently, stressed rats reduce their responding for food rewards [33, 34] and bias their decisions to low reward options due to an abnormal lack of motivation or anergia [33, 35]. Thus, both NE and DA can influence decisions under stress by affecting the decision strategy, motivation, and/or sensitivity to reward.

In the present study, we approach this topic with both pharmacology and fiber photometric analysis to elucidate the impact of stress, DA and NE on brain activity during decision making behavior. We induced stress pharmacologically with FG7142, a partial inverse GABAA agonist known to induce anxiety-related behavioral and physiological responses in a variety of species, including humans [36]. In a risky decision-making task, we show that a stress-induced bias towards safety shifted towards indifference by systemic administration of the α2A receptor antagonist, yohimbine. We further studied the temporal dynamics of NE and DA release related to aspects of trial outcome in the basolateral amygdala (BLA) and nucleus accumbens (NAc), respectively. In both structures, we found that pharmacologically induced stress modulated catecholamine release and that yohimbine led to opposing modulation.

## MATERIALS AND METHODS

Full details of materials and methods is provided in supplemental **Fig. S1.** All experimental procedures were approved by NIMH Institutional Animal Care and Use Committee, in accordance with the NIH guidelines for the use of animals.

### Subjects

Male Long-Evans rats (Inotiv, Indianapolis, IN, USA) were pair-housed in a temperature-controlled room (23.3 °C) under diurnal conditions (12:12 h light: dark). Rats were maintained at 90% of the free-feeding weight and water was available for at least 2hrs a day.

### Decision making behavior

Rats were tested on a decision making task using the operant touchscreen platform. Rats chose between two different stimuli presented on the left and right side of the touchscreen monitor (**Fig. 1A**). Nosepoke touches to the ‘safe’ stimulus (leaf) always resulted in the delivery of the small 50μl sucrose reward. Responses to the ‘risky’ stimulus (circles), delivered a small 10μl sucrose reward 75% of the time, or a large 170μl sucrose reward 25% of the time. Following stable baseline performance, rats (n=12) were treated IP with an adrenergic α2A-receptor agonist (guanfacine) and antagonist (yohimbine). Two weeks later, we induced stress with a pharmacological stressor, FG 7142, a GABAA inverse agonist **(Fig 2A)** which induces biochemical changes that mimic stress or anxiety similar to those elicited by mild aversive conditioning [37, 38], and therefore occurs independently of nociceptive stimuli. We then sought to alter the effects of the stressor by co-injecting it with the noradrenergic receptor specific drug.

**Figure 1.**
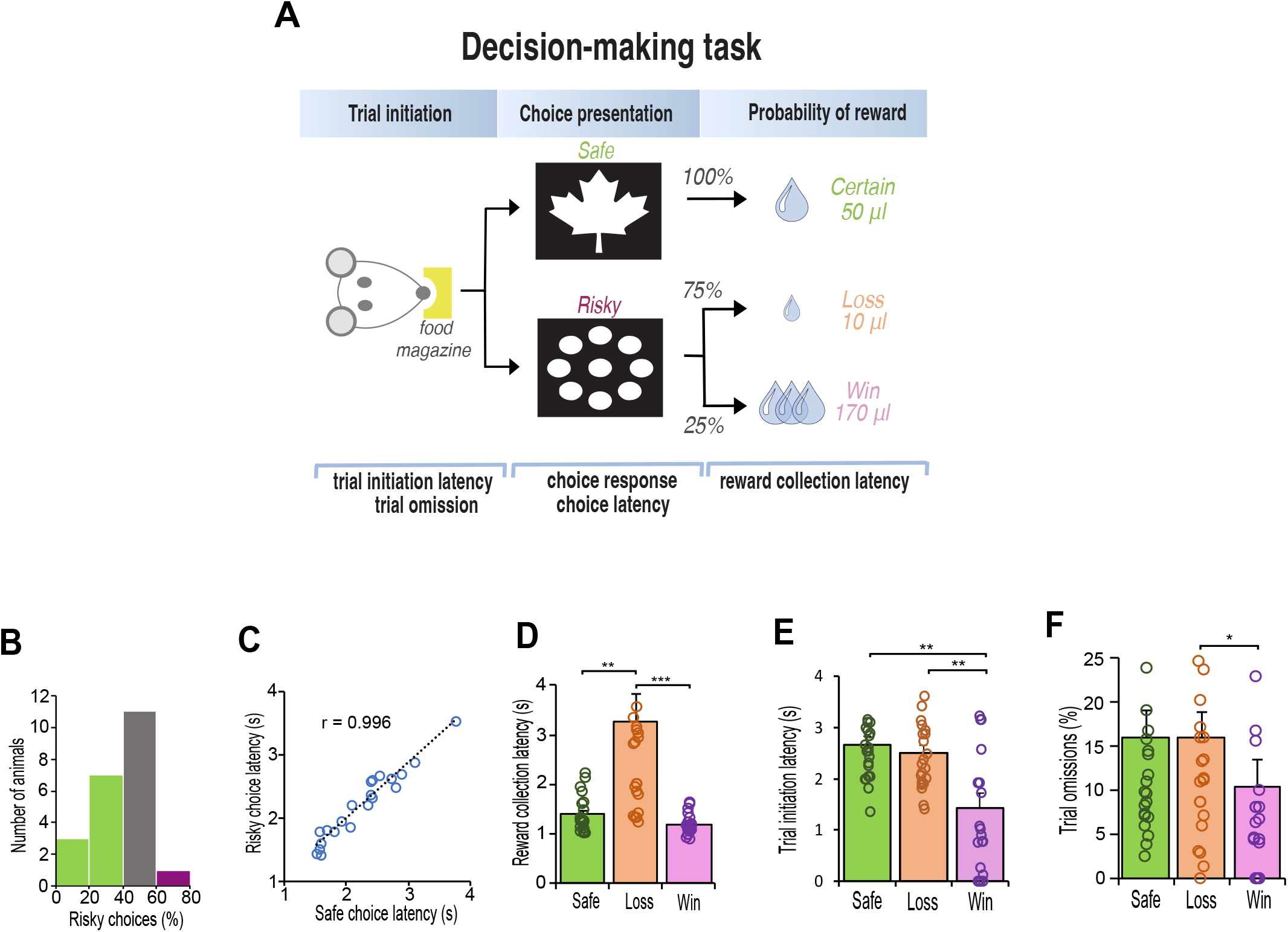
**Schematic illustration of decision-making task and baseline behavior after 15 days of training**. **A.** Rats initiated each trial with a nosepoke before choosing the ‘safe’ image (leaf) or the ‘risky’ image (circles). The safe image delivered a 50-μl reward. The risky image delivered a 10μl reward with 75% probability and a 170μl reward with 25% probability. The expected value of each choice was 50 μl of sucrose. **B**. Histogram of risk preference. Green indicates risk-averse rats (> 40% risky choices); grey indicates indifferent rats (> 40% and < 60%); purple indicates risk-seeking rats (> 60%). **C.** Latency to choose the safe or risky option were equal. **D**. Reward collection latency was slower for reward loss reward (10-μl sucrose) for all rats (n = 22, Friedman test, χ^2^ = 42.91, p < 0.001; Safe vs Loss, Adj. p = 0.002; Win vs Loss, Adj p < 0.001). **E**. Latency to initiate a trial was always faster after a win (170μl sucrose) relative to after a safe/certain reward or reward loss (n = 22, latency x reward outcome interaction F1,22 = 16.913, p > 0.001; Safe vs Loss, p = 0.001; Win vs Loss, p = 0.002). **F**. After losing a reward, rats were more likely to omit next trial than after winning a high reward (n = 22, Friedman test, χ^2^ = 7.747, p = 0.021; Win vs Loss, p = 0.031). Data are mean and S.E.M. *** p < 0.001, ** p < 0.01, * p < 0.05.

**Figure 2.**
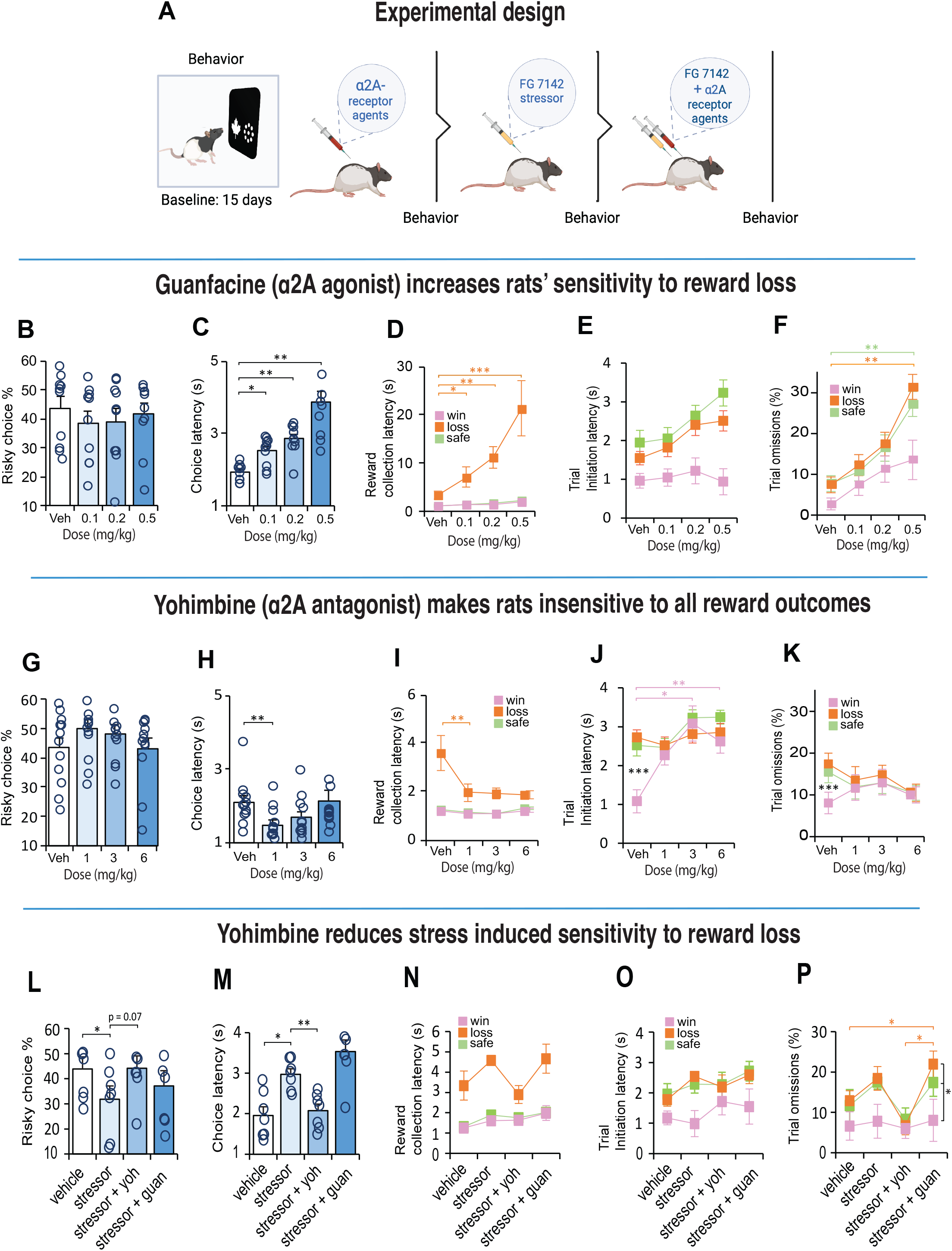
The effect of systemic α2A agonist and antagonists on decision making behavior. A. After 15 days of training, rats (n = 22) received a counterbalanced dose of an α2A agonist (guanfacine) and antagonist (yohimbine) 30 minutes before the test. Two weeks later, rats were injected with FG7142 to induce stress and tested again 30 minutes later for their decision making behavior. Subsequently, the animals were co-injected with the stressor and α2A receptor specific drug before behavioral testing. B, G. Risky choices are not affected by systemic α2A drug treatments. C, H. Guanfacine slowed down decision speed (Guanfacine, n =10, latency x dose interaction F3,27 = 24.512, p = 0.004; Veh vs all doses p < 0.05). In contrast, yohimbine made rats faster in their response speed (n = 12, Friedman test, χ2 = 15.5, p = 0.001; Veh vs Low, p = 0.01). D, I. Reward collection latencies increased with guanfacine (Guanfacine, n = 10, latency x dose interaction F3,23 = 14,41, p < 0.001; Veh vs all doses p < 0.05). Yohimbine made rats faster after loss (yohimbine, n = 10, latency x dose interaction F2,17 = 9.85, p = 0.002; Veh vs Low dose, p = 0.005). E, J. Trial initiation latency was generally faster after winning a high reward when injected with guanfacine, but not with yohimbine (Yohimbine, n = 12, latency x dose interaction F3,33 = 4.84, p < 0.001; Vehicle, Win vs Loss, p < 0.001). Yohimbine Low dose x latency interaction F2,22 = 0.457, p > 0.05. Low – low dose. Med – medium dose. High – high dose. Data are mean and S.E.M. *** p < 0.001, ** p < 0.01, * p < 0.05. L-P. Pharmacological stress induced by an i.p. injection of 4 mg/kg of FG 7142 reduced risky choices and increased their choice latency, while co-treatment of FG 7142 with yohimbine produced opposing effects; rats were faster in their choices and increased their choice of risky options to a level of indifference. Combined treatment of FG 7142 with guanfacine made rats slow and omit more trials after a reward loss.

### Fiber photometry

In a subset of pretrained animals (n = 9), we injected, unilaterally, a genetically encoded fluorescent NE sensor, GRABNE (pAAV-hSyn-GRAB_NE1m) in the basolateral amygdala (BLA), or a DA sensor, dLight in the nucleus accumbens (NAc) to monitor NE and DA neural activity while animals engaged in the decision-making task. Subsequently, we examined NE and DA responses following systemic injections of vehicle, yohimbine (1mg/kg) and FG7142 (4mg/kg).

### Statistical analysis

The behavioral data were processed using custom-written programs in R and analyzed using SPSS Statistics 25.0 (IBM, Chicago, IL, USA). Incomplete data with sessions comprising less than 30% of free choices trials were not used for statistical analysis. All data were tested for normality and transformed accordingly before statistical significance testing (see **Fig. S1** for full details).

## RESULTS

### High sensitivity to loss impacts motivational state

After 3 weeks of testing, rats displayed consistent choice preferences across three consecutive days (χ^2^ (2) = 0.302, p = 0.86). Similar to human choice behavior [40], the rats were either indifferent in their choices (n=11) or exhibited stable risk-aversion (n=10). Only one rat exhibited risk taking behavior by choosing the risky option more than 60% of the time **(Fig. 1B**). Regardless of their choice, the speed at which the animals made their response did not differ between the safe or risk option (t(21) = 0.97, p = 0.34), and correlated strongly (Pearson r(21) = .97, p < 0.001) such that the latency to choose the safe or risky option were equal (**Fig 1C**). However, their motivation to collect reward was highly influenced by their outcome, especially following selection of a risky option that led to a loss (χ^2^ (2) = 42.09, p < 0.001). Posthoc tests confirmed that the latency to collect reward was twice as fast following delivery of the safe-certain reward (Z = 3.47, p = 0.002) and a risk-win reward (Z = 6.48, p < 0.001) relative to a risk-loss (**Fig. 1D)**. Their motivation to initiate the next trial was also influenced by the outcome of the choice; after choosing the risky option that led to a reward loss, the speed to initiate the next trial was substantially slower than following a win (F (1, 23) = 16.91, p < 0.001, **Fig. 1E**), and in many cases resulted in trial omissions (Z = 2.56, p = 0.031, **Fig. 1F**).

### α2A – adrenoceptors modulate sensitivity associated with reward loss

We next examined how choice for safe or risky options were modulated by noradrenergic drugs that acted on the α2A-receptor. There is some evidence that stimulation of α2A-receptors affects some forms of decision making even in normal animals [41, 42]. To test this possibility, we first injected a cohort of trained rats with low, medium, and high doses (Fig. S1) of guanfacine (α2A- receptor agonist) and yohimbine (α2A-receptor antagonist), in separate sessions, each counterbalanced with vehicle **(Fig. 2A**). Neither drug had any impact on the animals’ preference for the choice at any dose (guanfacine: χ^2^ (3) = 4.09, p = 0.252; yohimbine: χ^2^ (3) = 5.5, p = 0.139; **Fig. 2B, G)**, but substantially altered the animals’ sensitivity to loss and motivational state, in opposite directions. In general, guanfacine demotivated the animals by slowing them down without impacting their motoric abilities. For example, their latencies to make a choice increased with higher doses (F(3, 27) = 24.512, p = 0.001; **Fig. 2C**), but their latencies to collect reward depended on the reward outcome (F(3, 23) = 14.41, p < 0.001). When the animal chose the risky option and experienced a reward loss, these animals were disproportionally slower in collecting the reward which got worse with increasing dose (F(3, 27) = 23.811, p < 0.001; **Fig. 2D).** The same animals, however, were distinctly fast to collect reward following a choice response that led to a win or a safe-certain reward at all doses. Thus, the long latencies could not be explained by mere sedation, but a specific sensitivity to reward loss. Similarly, the latency to initiate a trial following the different reward outcomes was impacted across all doses (F(3, 25) = 2.93, p = 0.056; **Fig. 2E)** as were the number of omission (F(6, 54) = 4.06, p = 0.002; **Fig, 2F**).

Opposite to the effects of guanfacine, antagonizing the α2A-receptors with yohimbine made the animals almost insensitive to the reward outcomes; their choice strategy remained constant (**Fig. 2G**), they were faster to make a choice response especially at the low dose (χ^2^ (3) = 15.5, p = 0.001; **Fig. 2H**) and comparatively faster than guanfacine (compare with **Fig. 2C**). A small difference emerged between reward loss and reward win collection latencies (F(2,18) = 7.75, p = 0.005), but relative to vehicle, the yohimbine made rats substantially faster to collect rewards even when the choice led to a large reward loss (**Fig. 2I**). Moreover, **Fig. 2J** shows that with yohimbine, the animals were so insensitive to the different reward values that their motivation to initiate the next trial was identical for all reward outcomes including after reward losses (F(2, 22) = 0.46, p = 0.64), but not with vehicle (F(2, 22) = 24.4, p < 0.001). A similar pattern was observed for omissions (Vehicle: F(2, 22) = 31.6, p < 0.001; yohimbine: F(2, 22) = 1.33, p = 0.29; **Fig. 2K**)

Since many psychiatric disorders characterized by risky decision making are associated with dysregulated dopamine transmission [43, 44], in a separate cohort of rats, we also examined the effects SKF 81297 (dopamine D1 receptor agonist) and SCH 23390 (dopamine D1 receptor antagonist) for comparison. The D1 antagonist, while not affecting the animal’s choice behavior affected the animals motivation by increasing their latencies following a reward loss, but there was no major impact on decision making behavior with the D1 agonist (**Fig. S2**).

### Yohimbine reduces stress-induced sensitivity to reward loss

We next examined if decision-making was sensitive to physiological stress. Due to the habituation caused by repeated exposure to stress, we gave rats a systemic 4mg/kg dose of a pharmacological stressor known as FG 7142 (henceforth known as stressor or FG stressor; **Fig 2A**). This drug is known to mimic the effects of uncontrollable stress linked to anxiety [45] and increases catecholamine turnover in various limbic associated areas including the BLA and NAc [46–48] thought to influence choice behavior in humans and animals [49]. The stressor veered rats’ choices towards safety (F(3, 18) = 6.408, p = 0.004; veh vs stressor, p = 0.016; **Fig. 2L**) and made them slower in their response (F(3, 18) = 25.859, p < 0.001; veh vs stressor, p = 0.026; **Fig. 2M**) suggesting they were less motivated to engage in risk taking behavior. Apart from an increased sensitivity to reward loss, other aspects of motivation were relatively intact (**Figs. 2N- P**). In humans, however, the co-administration of the stress hormone hydrocortisone with yohimbine diminishes loss aversion [11]. To test this possibility, we co-injected the FG stressor with yohimbine and found, that their concurrent actions tended toward reduced loss aversion in rats relative to the stressor alone (F(3, 18) = 6.408, p = 0.004; FG vs FG + Yoh, p = 0.07; **Fig. 2L**), and made them noticeably faster in their choices (p = 0.008; **Fig. 2M**). It also increased their motivation to collect low rewards, initiate trials and reduced the number of omissions after a reward loss (**Figs 2N-P**). In contrast, the behavior associated with combined guanfacine, and the FG stressor was equivalent to the stressor alone (all p>0.05, NS).

### FG stressor and yohimbine produce opposing effects on BLA-NE release

We next asked if the fluorogenic NE reporter GRABNE, could capture the rapid dynamic properties of NE release for decisions that led to different reward outcomes. We focused on the basolateral amygdala (BLA) because it is diffusely innervated by NE projecting neurons from the locus coeruleus [50–52], and NE levels in the BLA increase with presentation of stressful stimuli [24, 25]. We injected rats with an AAV that expressed GRABNE in the BLA and implanted an optic fiber above the injection site (**Figs. 3A, S3A**). After a minimum of 4 weeks to allow the expression of the viral sensor, rats were placed in the test chambers and assessed on their decision-making while recording the GRABNE fluorescence. The fluorescent signal was aligned to four specific events in the trial: the choice response, reward collection, before trial initiation and after trial initiation (**Fig. 3B)**. Changes in NE release were not observed in the BLA during the decision itself such that the kinetics of the NE response were relatively equivalent before and after the choice (**Fig. 3C**). A double activation of signal for NE release in the BLA was observed just before and immediately after reward collection regardless of the trial outcome (**Figs. 3D, S4A-C).**

**Figure 3.**
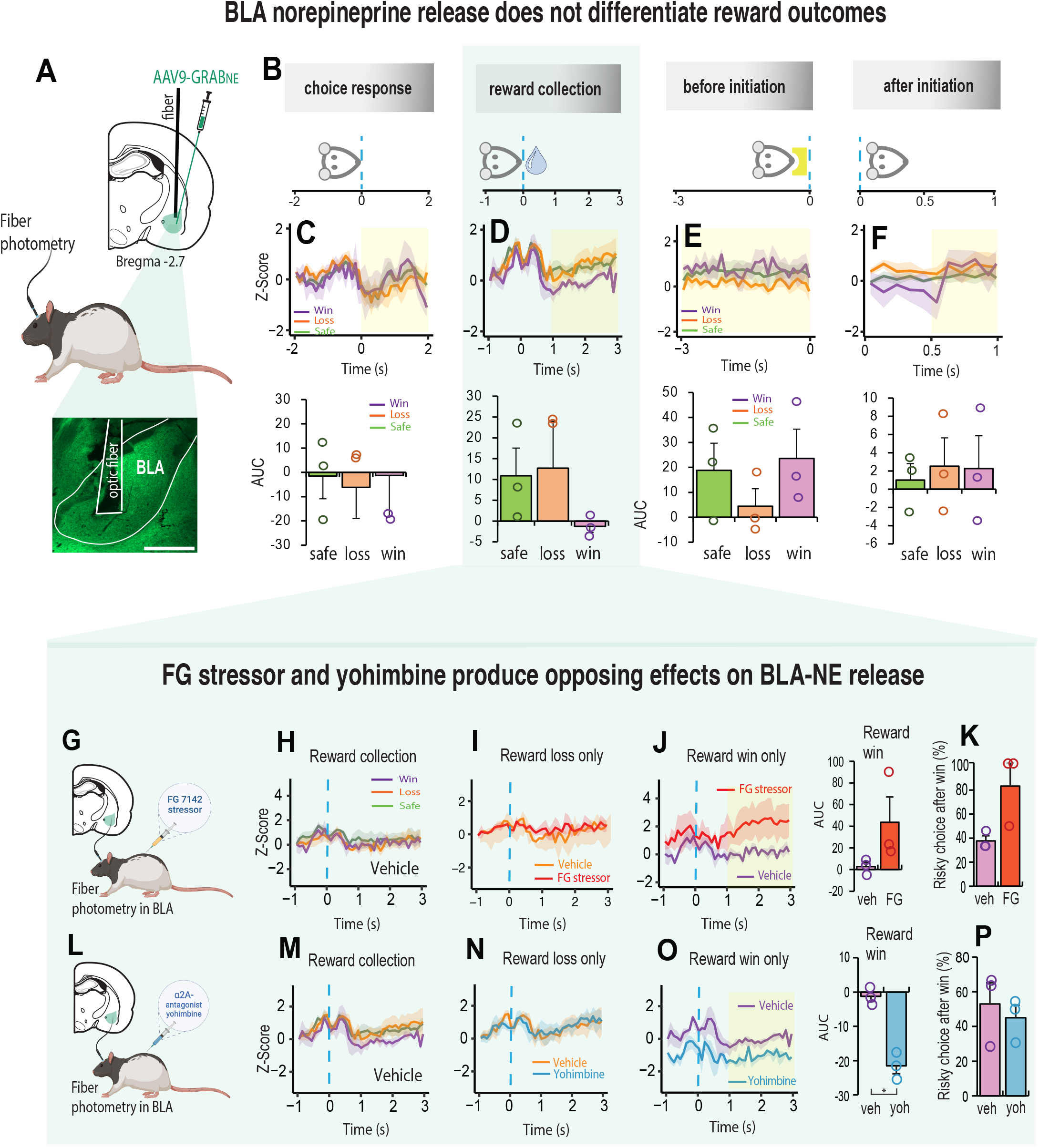
FG stressor and yohimbine produce opposing effects on BLA-NE release A. Schematic illustration of AVV9-GRAB_NE1m expression in the BLA and histology showing optic fiber placement above the injection site. B. Schematic illustration of how NE signal was aligned to four specific events in the trial. Choice response: the rat touched the screen and moved towards reward collection. Reward collection: rat entered the food magazine to collect the reward. Before initiation: 3 second before the next trial was initiated. After initiation: 1 second after trial initiation, during which rat moves towards touchscreen for the next choice. C-F Top row. Mean noradrenaline signal during choice, reward collection, after initiation and before initiation (n = 3). The dynamics of NE release are different for reward wins (purple), safe/certain rewards (green) and reward loss (orange). The yellow shaded area assigns the period used for the area under the curve analysis (AUC). C-F Bottom row. Shaded yellow area assigns the period for the area under the curve analysis (AUC). G. Schematic illustration of fiber photometry probe in BLA combined with FG7142 injection. H. NE signal in the BLA aligned to reward collection for vehicle (n = 3). I. NE signal in BLA following vehicle (orange) and FG7142 (red) treatment are indistinguishable reward collection resulted in reward loss. J. FG stress (red) induced NE signal for collection of high reward win increased relative to vehicle (purple). K. The increase in NE signal after collecting a high reward was associated with a higher propensity to make risky choices. L. Schematic illustration of fiber photometry probe in BLA combined with yohimbine injection. M. NE signal in the BLA aligned to reward collection with vehicle. NE signal did not differentiate between win (purple), safe (green) and loss (orange) reward outcomes. N. NE signal for reward collection after a reward loss was indistinguishable between vehicle (orange) and yohimbine (blue). O. NE signal in BLA for collection of a high reward win in yohimbine (blue) treated rats reduced relative to vehicle (purple). P. The reduction in NE signal did not influence the subsequent choice. Data are mean and S.E.M. AUC – area under the curve. Veh – vehicle. FG – FG7142. Yoh – yohimbine. Scale bar 500 µm. ** p < 0.01, * p < 0.05.

Subsequently the signal declined especially for choices that led to large reward wins. The signal for the remaining trial events did not differentiate the reward outcomes **(Figs. 3E, F)**. Thus, NE release in the BLA does not correlate directly with choices influenced by risk or uncertainty. However, since yohimbine and the FG stressor altered the animal’s sensitivity to reward outcome when injected systemically (**Figs. 2L-P**), we measured NE signal in the BLA following the injection of each of these agents (**Figs. 3G, L)**. We found these drugs to have opposing effects on NE release but only when we aligned the signal to reward collection **(Figs. 3G-P)**. While the stressor had no impact on the NE response for reward losses (**Fig. 3I**), it increased the NE signal following a large reward win (**Fig. 3J**; t2 = 2.02, p = 0.181). Although it did not affect decision strategy (t2 = 2.675, p = 0.11), there was a higher propensity to make risky choices after a reward win **Fig. 3K**). Conversely, systemic yohimbine reduced NE release in the BLA relative to vehicle when the choice led to a large reward win (t2 = 9.407, p = 0.011) but this did not influence the subsequent choice (**Fig. 3L-P**). The BLA-NE signal did not change for either drug when the reward collection followed a loss (**Fig. 3I, N**). These results suggest that α2A – adrenoceptors can potentially suppress or shift the effects of stress induced risky behavior by making the animal choose safer options.

### Stress induced DA response in the NAc is modulated by α2A – adrenoceptors

We also asked if stress induced changes in motivation could be differentially modulated by dopamine (DA). There is much evidence however, that stress has profound effects on the mesoaccumbens dopamine system [30, 31, 53] and that projections from dopaminergic nuclei to the nucleus accumbens (NAc) play an important role in motivated decision-making behavior [54, 55]. Accordingly, we first recorded changes in fluorescence of the genetically encoded dopamine (DA) sensor, dLight expressed in the NAc to examine changes in the DA response during decision-making (**Fig. 4A, S3B**). The highest peak of DA release in the NAc was observed after the choice was made, and it related to the value of the future reward (**Figs. 4B-C, S4D-F**). Thus, DA release was high when there was an increase in future reward value (a reward win), inhibited when there was decrease in future reward value (a reward loss), and intermediate when the future reward was low but certain (Choice: F (2, 10) = 57.39, p < 0.001; **Fig. 4C**). Moreover, the DA response for the winning choice peaked high during reward collection and elevated again *after* reward collection (F (1, 5) = 6.95, p = 0.031; **Fig. 4D**). High DA release after a win persisted to some degree until the animal initiated the next trial (before initiation: F (1, 5) = 3.3, p = 0.079; **Fig. 4E),** which could potentially explain the high motivational state characterized by faster trial initiations and reduced omissions (**Fig 1D** **- F**). After the next trial was initiated, the DA signal reversed; the previous reward wining outcome resulted in a reduction in DA release whilst the previous low reward outcomes enhanced DA release in the NAc (after initiation: F (2, 10) = 8.06, p = 0.008; **Fig. 4F**).

**Figure 4.**
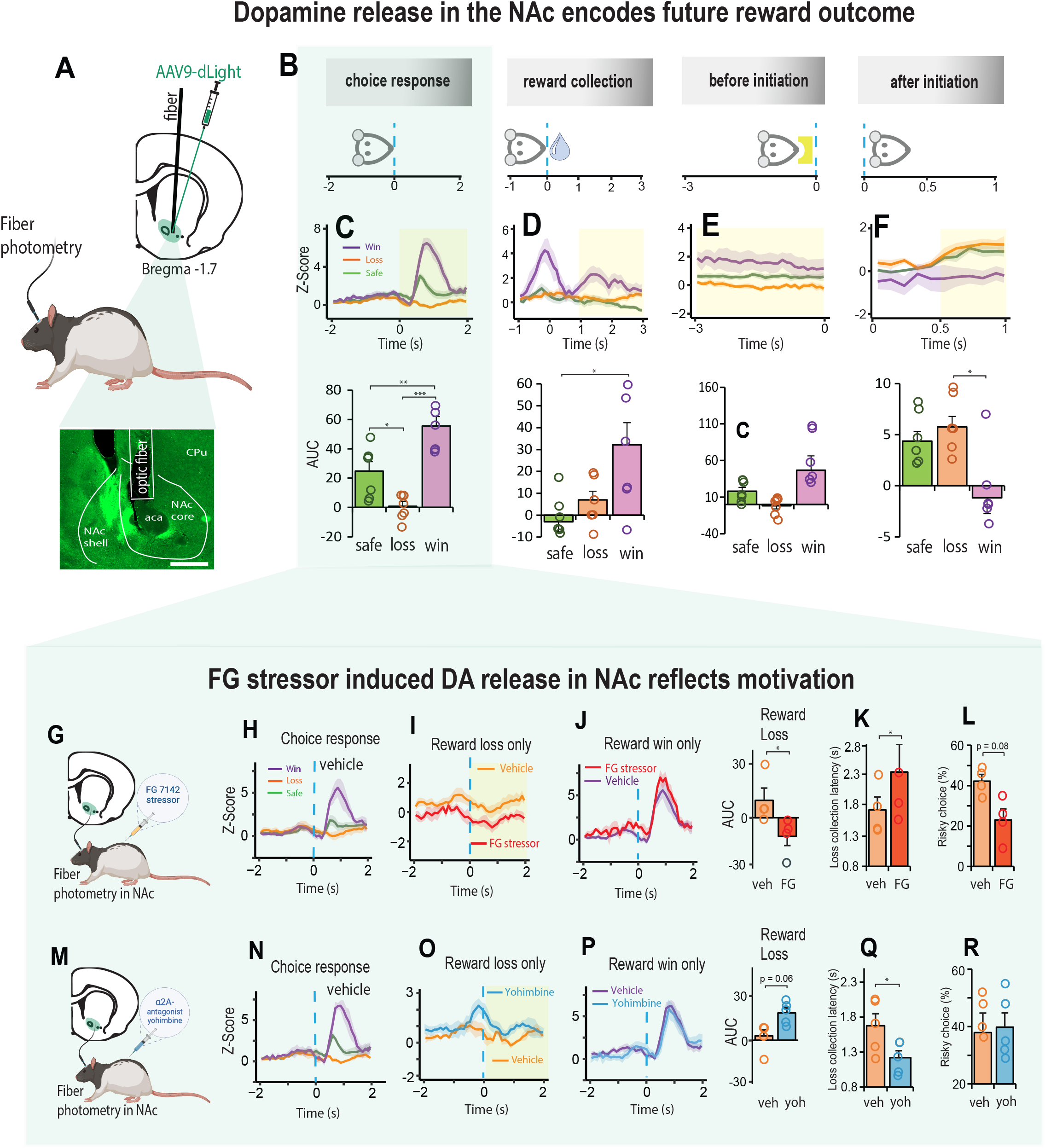
**Stress induced DA release in the NAc reflects motivation during decision-making**. **A** Schematic representation of an AVV9-dLight expression in the NAc and histology showing optic fiber placement above the injection site. **B**. Schematic illustrating how NE signal was aligned to four specific events in the trial. Choice response: the rat touched the screen and moved towards reward collection. Reward collection: rat entered the food magazine to collect the reward. Before initiation: 3 second before the next trial was initiated. After initiation: 1 second after trial initiation, during which rat moves towards touchscreen for the next choice. **C - F.** DA signal in NAc during the four trial events shown in B (n = 5). **C.** Strength of DA release aligned to the choice response varied systemically as a function of expected reward value such that DA release was high when there was an increase in future reward value (a reward win: purple), inhibited when there was decrease in future reward value (a reward loss: orange), and intermediate when the future reward was low but certain (green). Win vs Loss, p < 0.001; Win vs Safe, p = 0.002; Loss vs Safe, p = 0.017). Shaded yellow area assigns the period for the area under the curve analysis (AUC). **D.** The dynamics of the DA signal was greater after collection of a large reward win than after safe/loss reward collection. Win vs Loss, p = 0.097; Win vs Safe, p = 0.029). **E.** Before initiating the next trial, the DA signal was slightly elevated if the previous choice was a win. **F.** The DA signal lowered substantially once the next trial was initiated if the previous choice was a win. Win vs Loss, p = 0.024; Win vs Safe, p = 0.054). **G.** Schematic representation of dopamine signal collection in NAc combined with FG7142 injection. **H.** DA signal during aligned to choice response (n = 4). **I.** Reduced DA signal when choice response led to a reward loss for Veh (orange) and FG stressor (red). **J.** Elevated DA signal for choice response leading to future reward win with stressor (red) and vehicle (purple) were similar. **K-L.** Reduced DA signal with stressor injection leading to reward loss was associated with long reward collection latencies and an increase in choices towards safety. **M.** Schematic representation of DA signal in NAc combined with yohimbine injection. **N.** DA signal aligned to the choice phase following Veh injection (n = 5). **O.** DA signal when choice response led to reward loss with yohimbine (blue) was higher relative to vehicle (orange). **P.** DA signal when choice response led to reward win following vehicle (purple) and yohimbine (blue) were indistinguishable. **Q-R.** Decrease in DA signal following reward loss was associated with faster reward collection latencies but did not affect the animal decision strategy (loss collection latency, t4 = 2.783, p = 0.049). Data are mean and S.E.M. AUC – area under the curve. Veh – vehicle. FG – FG7142. Yoh – yohimbine. Scale bar 500 µm. *** p < 0.001, ** p < 0.01, * p < 0.05.

We next examined changes in DA release in the NAc following systemic injections of the FG stressor (**Fig. 4G-L**) and found that it had no impact on the DA signal for choices that led to a reward win, but for those choices that led to a reward loss, the DA signal declined relative to vehicle **(Fig. I-J).** Moreover, as we observed through systemic injections (**Fig. 2L**), there was an associated increase in loss sensitivity characterized by slower collection of low rewards and increased aversion to risk (AUC, t3 = 4.299, p = 0.023; loss collection latency, t3 = 3.875, p = 0.03; % risky choice, t3 = 2.6, p = 0.08; **Fig. 4K, L)**. We then discovered that the FG stressor and yohimbine had opposing effects on NAc-DA raising the possibility that the sensitivity associated with loss could be potentially reduced with α2A - adrenoceptors (**Fig. 4M-R**). This not only increased DA release in response to reward loss (**Fig. 4O**), it increased the rat’s motivation by making them faster to collect a low reward without affecting their decision strategy (**Fig. 4Q, R**; AUC, t4 = 2.55, p = 0.063; loss collection latency, t4 = 2.783, p = 0.049).

## DISCUSSION

There were three main findings: 1) pharmacological stress shifted behavior toward safe decisions, and this effect was reduced by yohimbine, 2) following large gains after a risky choice, the level of NE release in the BLA was increased by a pharmacological stressor but decreased by yohimbine, 3) following a loss after a risky choice, the level of DA in the NAc was decreased by a pharmacological stressor but increased by yohimbine. These opposing effects of the FG stressor and yohimbine on catecholamine release in these critical structures during the outcome of risky decisions provides a plausible explanation of why yohimbine shifts choice behavior in the face of a pharmacological stressor. Specifically, it points to the role of α2A – adrenoceptors as being critical to the remediation of the stress effects.

### Opposing influence of α2A adrenoceptor activation

One major finding of our study is that NE modulation at α2A adrenoceptors has a powerful influence on how reward loss is incorporated into the animals’ decision. We found that, similar to the pharmacological stressor, FG 7142, loss aversion was disproportionately high in animals injected with the α2A agonist, guanfacine; they were slow in making choices, collecting their rewards, and initiating trials, but only when their choice resulted in a low reward outcome. When the choice outcome resulted in a high reward, rats showed normal levels of motivation and speed of response even at the high dose, confirming that the demotivating effects of systemic guanfacine following reward loss were not due to sedative effects of the drug. Guanfacine, through its postsynaptic actions in the prefrontal cortex, is known to enhance cognitive functions that subserve attention and working memory [56, 57]. One possibility is that while guanfacine made the α2A adrenoceptor highly sensitive to decisions that lead to major losses, it functioned to enhance or focus attention most efficiently towards choice outcomes with positive consequences.

In contrast, when the α2A receptors were antagonized with yohimbine, these same animals showed exaggerated focus and speed, with no evidence of loss aversion. In fact, in their eagerness to respond, they often failed to discriminate the different reward values associated with the choice outcome. Consequently, these animals worked equally fast for all trials even when their chosen option lead to a loss. While it is possible that the fast, indiscriminate responding with yohimbine was caused by an impulsive-like state [58–60], they were not prone to make riskier choices. In fact, despite the robust alterations in the animals’ motivational state, neither yohimbine nor guanfacine, when systemically administered alone, had any impact on the animal’s decision strategy.

### α2A adrenoceptor modulation of stress induced catecholamine release

Although systemic injections of the FG stressor shifted rats’ preferences for low-risk choices, its impact on NE activity in the BLA was markedly different. The photometry trace revealed an elevated NE signal in BLA neurons for the large reward win, which further inclined the rat to make high-risk choices. The high NE signal is consistent with the general finding that stressful stimuli increase NE release in the BLA [24, 48, 61]. In fact, when rats were not stressed, there was very little variation in NE release in the BLA suggesting that basal levels of BLA-NE have little impact on decision-making under risk. Notably, the NE signal did not change when choices resulted in a reward loss suggesting that NE in the BLA, while sensitive to emotionally stressful or fearful contexts [62, 63], may not encode the affective consequence of loss, or ‘loss aversion.’ This difference may be due to the nature of the stressor or its intensity which differentially impacts the activity of noradrenergic neurons in different brain regions [64–66]. Since benzodiazepine receptor binding is altered by stress [67, 68], and NE acts on GABAergic cell populations indigenous to the BLA region [69, 70], the inclination to be risky may be related to impaired NE modulation of GABA transmission in the BLA. Our finding that systemic injections of yohimbine reduced the BLA-NE signal without affecting decision strategy is consistent with this hypothesis.

We note that unlike other published reports [71, 72], the effects of the DA D1 agonist/antagonist on decision making were modest at best (Fig. S2), but not uncommon [73–75]. In our case, the DA D1 agonist and antagonist had a noticeable effect on motivation which was further supported with the photometric analysis of DA release in the NAc. In the absence of stress, the DA signal in the NAc encoded both, the reward value associated with the choice, as well as the magnitude of the high reward which was sustained for some time after reward collection. Notably, the size of the NAc-DA signal positively correlated with the rat’s motivation (**Fig. S5**), a finding contrary to that of Eshel et al., [54] where motivation, characterized by the willingness to overcome the cost of working in mice, was found to be negatively correlated with NAc-DA activity. In the present study, the sensitivity to reward loss exacerbated when the animal was stressed with FG 7142. This resulted in a reduced DA signal in the NAc, whereas yohimbine increased the DA signal. In fact, with yohimbine, the NAc-DA signal was high even *before* the choice was made, and this predicted the animals increased motivation to decide faster on their options and collect their rewards, even for low rewards. Thus, although DA activity in the NAc is sensitive to experienced loss during stress, we found that the reduced motivation exhibited by these animals for low reward choices can be potentially countered with an α2A adrenoreceptor interaction with GABAA receptors in the NAc.

### Concluding remarks

We applied methods of behavior, psychopharmacology and fiber photometry to measure the neural dynamics of stress modulators in brain regions that affect decision-making in rats. Like humans, we showed that rats imitate the loss aversion effect and its associated outcome-based motivations. Our results support the proposal that stress induced changes in catecholamine release in the BLA and NAc can directly influence loss sensitivity, choices and motivation, which can be modulated by the α2A adrenoreceptor antagonist, yohimbine. Stress associated catecholamine release exacerbates vulnerability to a variety of clinical conditions in which patients engage in decision making involving risks and rewards. Elucidating the complex interplay between neuromodulatory circuits that mediate this form of decision making will improve our understanding of the dysfunctional pathways linking the stress response to suboptimal life-altering choices.

## ACKNOWLEDGMENTS

We thank Dr. Gloria Laryea for initial discussion of task design. We also thank Dr. Sean Bradley, Ms. Alice Graham and Mr. Kevin Cravedi in the NIMH Rodent Behavioral Core for resources and support for rodent behavioral testing. CJT is now at Georgetown University, DC, USA. AA is now a military contractor working with soldiers on their decision-making and attention control in stressful situations.

## AUTHOR CONTRIBUTIONS

Conceptualization, design and methodology VV, YC; investigation and data collection VV, CJT, AA, MHL, FM; Formal analysis VV; Figures and visualization VV, YC; resources YC; Writing original draft VV, YC; writing review and editing VV, CJT, FM, MHL, YC.

## Supplemental Information

### Figure S1: Materials and Methods Subjects

Male Long-Evans rats (Inotiv, Indianapolis, IN, USA) weighing 250-280g at the start of behavioral training were used for these studies. They were pair-housed in a temperature-controlled room (23.3 °C) under diurnal conditions (12:12 h light: dark). All testing occurred at a regular time during the light period. Rats were maintained at 90% of the free-feeding weight and water was available for at least 2hrs a day. All experimental procedures were approved by NIMH Institutional Animal Care and Use Committee, in accordance with the NIH guidelines for the use of animals.

### Decision-making behavior

#### Touchscreen Operant Platform

Behavioral testing was in eight automated touchscreen operant chambers (Lafayette Instrument Company, Lafayette, IN, USA) each comprising a standard operant chamber fitted with a touchscreen. Each chamber was individually housed in a sound- attenuating cabinet and illuminated by a 3W houselight mounted on the ceiling of the cabinet. Computer graphic stimuli composed of white geometric symbols on a black background were presented on a touch-sensitive monitor (9” W × 10” H). A black mask made of anodized aluminum was attached to the face of the screen approximately 1.5cm from the surface of the display. The mask served to restrict the rats access to the screen except through two response windows (3” W × 3” H). Opposite the touchscreen was a precision liquid food pump (Lafayette Instrument Company, Lafayette, IN, USA) which delivered 10% sucrose solution into a food magazine. A light-emitting diode illuminated the food magazine. Magazine entries were detected by photocells located at the entrance of the food magazine. The apparatus and online data collection for each chamber were controlled with ABET II Software for operant control (Lafayette Instrument Company, Lafayette, IN, USA) interfaced with the Whisker control system for research (Cardinal and Aitken, 2010).

#### Pretraining

Following habituation to the testing chamber, rats were trained to enter the food magazine to collect to 50 μl sucrose reward. When rats were able to make 50 food magazine entries and thereby retrieve 50 rewards, they were trained to retrieve 50 rewards by touching one of two illuminated white squares (3”x 3”) presented on the touchscreen. Next, the rats were trained to initiate a trial by making a nose poke entry into an illuminated food magazine which resulted in the presentation of two white squares. In the first phase, a nosepoke touch to either of the white squares led to sucrose delivery. In the second phase, only one square was presented on either the left or the right side of the screen. A minimum of 50 nosepoke touch responses were required for each phase. Each pretraining session lasted 30 mins. In the last phase, the nose poke to receptacle initiated a white square on one of the randomized sides. To pass the training, rats were expected to touch each side more than 50 times. Once all 4 training phases were completed, rats were exposed to the behavioral task described in Fig. 1A. On average, rats were pretrained for ∼ 5 days. Following habituation to the chamber, rats were trained to reliably initiate trials, touch the screen and collect 10% sucrose solution as a reward.

#### Behavior

Rats chose between two different computer graphic stimuli presented on the left and right side of the touchscreen monitor (Fig. 1A). Each stimulus indicated differences in reward size and probability of outcome. Responses to the ‘safe’ stimulus (leaf) always resulted in the delivery of the small 50μl sucrose reward. Responses to the ‘risky’ stimulus (circles), delivered a small 10μl sucrose reward 75% of the time, or a large 170μl sucrose reward 25% of the time. The left/right positions of the risky and safe images were pseudorandomly determined thereby eliminating the potential confound of a side bias. Importantly, the expected value of the reward remained the same regardless of the animal’s choice. Each session started with 50 forced trials during which the safe or risky stimulus was presented to demonstrate the outcome associated with the stimulus. The remaining 200 trials were free choice trials in which rats could choose between both stimuli.

Each trial was signaled by the illumination of the magazine and house light. A nosepoke entry into the food magazine triggered the presentation of the stimuli for 10 sec. Following a successful response, the stimuli disappeared, all lights extinguished, and the chamber entered an intertrial interval (ITI) state of 10 sec. The next trial was signaled by the illumination of the food magazine and the houselight. Failure to make a nose poke entry with the 10 sec stimulus duration was recorded as an omission, and the box was returned to the intertrial state. The trial was then repeated until the rat received the full complement of 200 forced choice trials for that session.

Rats reached stable baseline performance in three weeks. Their fraction of risky choices across three consecutive days varied by less than 15 % (mean range = 7 %, and s.d. = 3). Rats that showed persistent side biases were excluded (n=2). The following formula was used to calculate the percentage of risky choices: % Risky = (Number of risky choices (Losses and Wins)/Total number of choices) *100). We also reported: choice response latency (time between trial initiation and topuch response), reward collection latency (time after choice response and collection of reward type (i.e., safe, loss or win), proportion of omission (failure to initiate the next trial after reward collection). The following formula was used to calculate the proportion of omission: (% omissions = (N of omissions after particular reward type)/(N of omissions and responses after particular reward type)

### Systemic pharmacology: Drug preparation and experimental design

All drugs were administered systemically i.p. and counterbalanced with vehicle. Doses of drugs were calculated as the salt and dissolved in the appropriate vehicle. **Table 1** lists all drugs, dosages, and dissolving vehicle. Behavioral testing occurred 30 minutes after the injection. Drug test days were followed by a drug free day of no testing. Animals were then tested on the baseline schedule until performance stabilized before the next treatment. All drugs were purchased from Tocris (Tocris Cookson Inc., Ellisville, Missouri, USA). Dimethyl sulfoxide (DMSO) and 2- hydroxypropyl-β-cyclodextrin (HBC) as dissolving agents were purchased from Sigma-Aldrich (St. Louis, MO, USA).

**Table 1.**
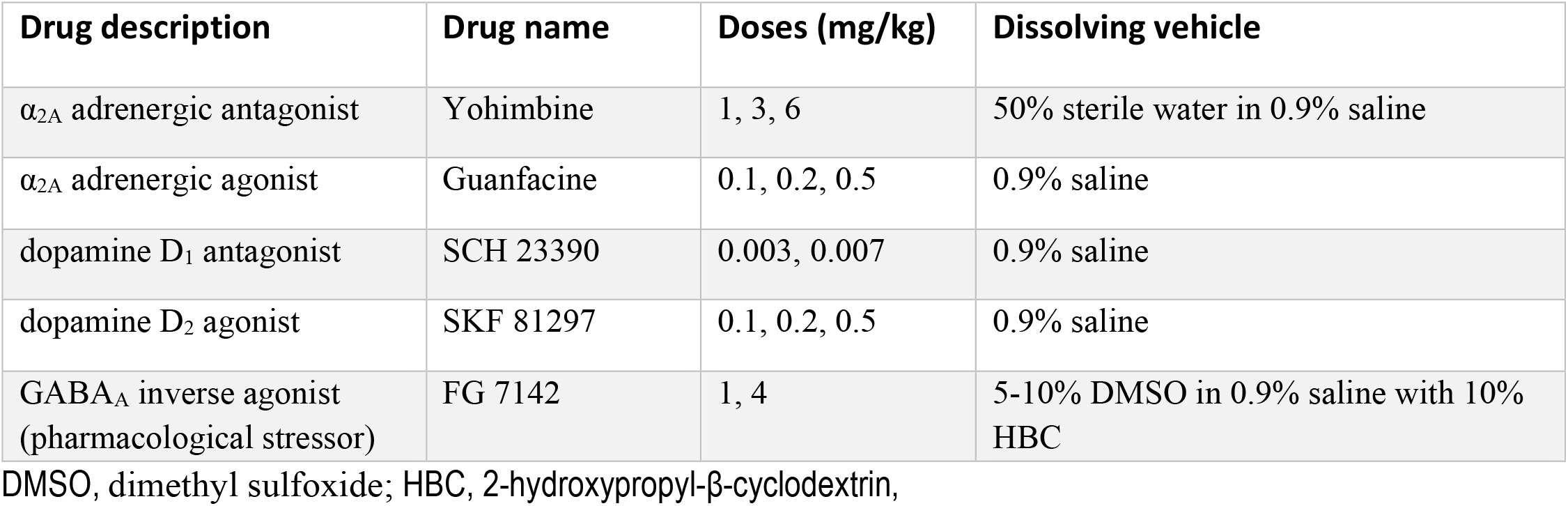
Summary details of drugs used in study.

| Drug description | Drug name | Doses (mg/kg) | Dissolving vehicle |
| --- | --- | --- | --- |
| $\alpha_{2A}$ adrenergic antagonist | Yohimbine | 1, 3, 6 | 50% sterile water in 0.9% saline |
| $\alpha_{2A}$ adrenergic agonist | Guanfacine | 0.1, 0.2, 0.5 | 0.9% saline |
| dopamine $D_1$ antagonist | SCH 23390 | 0.003, 0.007 | 0.9% saline |
| dopamine $D_2$ agonist | SKF 81297 | 0.1, 0.2, 0.5 | 0.9% saline |
| GABA<sub>A</sub> inverse agonist<br>(pharmacological stressor) | FG 7142 | 1, 4 | 5-10% DMSO in 0.9% saline with 10% HBC |

DMSO, dimethyl sulfoxide; HBC, 2-hydroxypropyl- β-cyclodextrin,

Following stable baseline performance, we examined the animals’ choice of risk and safe options following changes to dopaminergic or adrenergic receptor activity (Fig. 2A). Rats (n = 12) received injections of an adrenergic α2A-receptor agonist (guanfacine) and antagonist (yohimbine). Two weeks later, we induced stress in all animals by injecting them with a pharmacological stressor, FG 7142. We then sought to alter the effects of the stressor by co-injecting it with the noradrenergic receptor specific drug. Since stress is also associated with dopamine (DA) release in various brain regions, for comparison, we repeated the procedure in another cohort of rats (n = 10) who received injections of a DA D1 receptor agonist (SCH 23390) and antagonist (SKF 81297).

### Fiber photometry: Viral injection and fiber implants

In a subset of pretrained animals (n = 9), we injected, unilaterally, a genetically encoded fluorescent NE sensor, GRABNE (pAAV-hSyn-GRAB_NE1m; Addgene #123308, gift from Yulong Li) into the basolateral amygdala (BLA), and a DA sensor, dLight (pAAV-syn-dLight1.3b; Addgene #135762, gift from Lin Tian) into the nucleus accumbens (NAc), to monitor NE and DA neural activity during stress induced decision-making. For all procedures involving local injections of injection and probe implantation, rats were anesthetized with isoflurane gas (5% induction, 2% maintenance) and secured in stereotaxic headholder (David Kopf Instruments, Tujanga, CA, USA). The scalp was retracted to expose the skull and craniotomies were made directly above the BLA (A/P -2.7 mm, M/L 4.9 mm, D/V -7.6) and NAc (A/P 1.7 mm, M/L 1.6 mm, D/V -7.3 mm).

Viral injections were made using a pulled glass micropipette (WPI, USA) controlled by a Nanoliter 2020 injector (volume 300 nl at a rate of 100 nl/min). The virus was allowed to diffuse for 10 min before a slow withdrawal. Fiber optic cannulas (NA 0.66, 400-μm core diameter) were implanted

0.1 mm dorsal to the viral injection site (Doric Lenses, Canada). Cannulas were affixed with dental cement and stainless sterile screws to secure them in place.

### Fiber photometry recordings

Following a minimum of two-weeks after surgery, rats were re-trained to acquire stable decision-making performance (∼ 4 weeks). We first monitored NE and DA activity in the BLA and NAc, respectively while animals engaged in the decision-making task. Subsequently, we examined NE and DA responses following systemic injections of vehicle, yohimbine (1mg/kg) and FG7142 (4mg/kg).

Fiber photometry data were acquired with the RZ10X processor integrated with software Synapse v.96 (Tucker-Davis Technologies, Inc. USA). Light emitted from LED drivers integrated in the system (465 nm modulated at 330 Hz to excite dLight and GRAB-NE, and 405 nm modulated at 210 Hz for the isosbestic control) were transmitted through a Mini-cube fiber photometry apparatus (Doric Lenses) and low-autofluorescence patch-cord (400 μm core, 0.57 NA) connected to the implanted fiber-optic cannulas via a pigtailed rotary joint (Doric Lenses). The emitted signals were sent back to the Mini cube for filtration and detection by the integrated photosensors and demodulated in the Synapse software. In parallel, the RZ10X processor received time stamps of the behavioral events through a TTL breakout adapter (Lafayette Instruments, IN, USA). Raw fluorescence signals with behavioral time stamps were extracted into the Fiber photometry Modular Analysis Tool (pMAT) for further analysis [39]. Raw dLight or GRAB_NE signals were normalized to the isosbestic signal and transformed into delta F/F values. A custom-made R code and pMAT were used to calculate the delta F/F, Z-score and area under the curve (AUC) values used for analyses.

### Verification of fiber placement and viral expression

Rats were perfused transcardially, with a working solution of phosphate buffer saline (1X PBS) followed by 4% paraformaldehyde (PFA) dissolved in PBS. The brains were extracted and post- fixed in 4% PFA overnight at 4 °C, and then dehydrated in 30% sucrose in PBS for a week. The brains were then cryo-sectioned to 40 µm thickness using a freezing microtome (Leica Biosystems, USA). Sections were mounted on glass slides with Vectashield antifade mounting medium (Vector Laboratories, USA). High resolution images were taken with a microscope scanner (Axio Scan 7; Zeiss). Animals with misplaced cannulas or viral expression were excluded from analysis.

### Statistical analysis

The behavioral data were processed using custom-written programs in R and analyzed using SPSS Statistics 25.0 (IBM, Chicago, IL, USA). Incomplete data with sessions comprising less than 30% of free choices trials were not used for statistical analysis. All data were tested for normality and transformed accordingly before statistical significance testing. For comparisons between two groups *t*-tests were used. In cases when the data did not fit the assumptions of the test, the non- parametric Mann–Whitney or Wilcoxon matched-pairs tests were used. For repeated measures ANOVA, data was assessed for homogeneity of variance using Mauchly’s sphericity test. When this requirement was violated for a repeated measures design, the F term was tested against degrees of freedom corrected by Greenhouse–Geisser to provide a more conservative p value for each F ratio. Otherwise, nonparametric Friedman’s test (χ^2^) was applied with differences compared with posthoc Wilcoxon signed-rank tests (Z) adjusted with a Bonferroni correction. Pearson correlation (r) was used to describe the linear relationship between two correlated variables.

---

**Figure S1.** Methods and Materials. Detailed account of apparatus, behavioral procedure, drug preparation and experimental design, and statistics.

**Figure S2.**
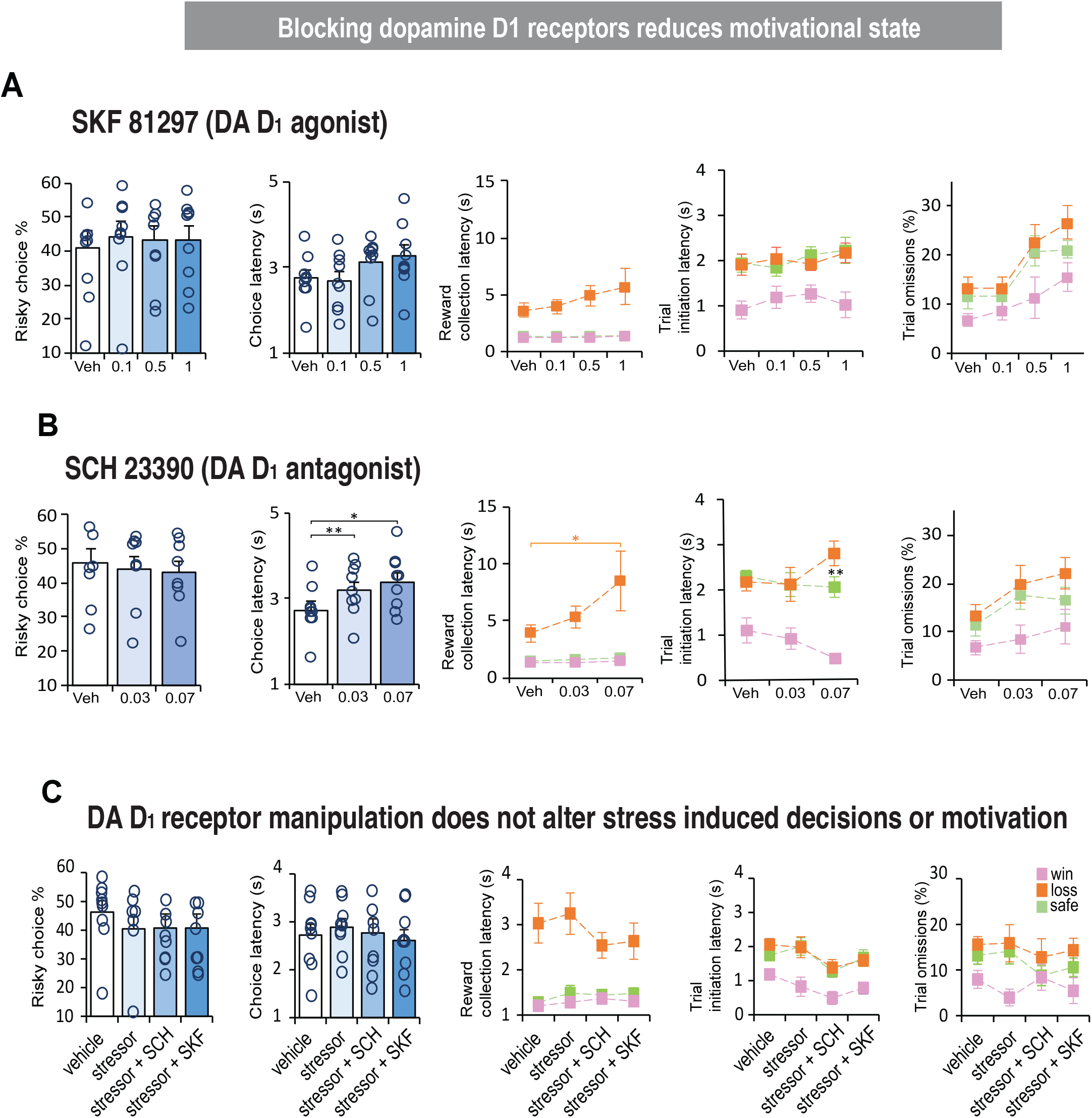
Blocking dopamine D1 receptors reduces motivational state. A. Systemic injections of the D1 agonist had no major impact on decision making behavior in that aspects of motivation including speed of response and reward collection following safe or risky choices were within the normal range. B. In contrast, the D1 antagonist, while not affecting the animal’s choice behavior, did alter their motivation, which in some way was similar to the effects of the α2A- receptor agonist, guanfacine (Fig. 2B). First, these animals were slow in their choice response for both doses relative to vehicle (0.03 mg/kg, p = 0.002; .07 mg/kg, p = 0.024). Second, there was a dose dependent increase in reward collection latency but only following a choice that led to a reward loss (F(2,13) = 5.40, p = 0.021). As expected, motivation for initiating a trial in this cohort of animals was relatively fast following a win, especially at the highest dose (0.07 mg/kg; F(2,16) = 50.312, p > 0.001). C. We also asked if stress induced changes in motivation could be differentially modulated by. We first co-injected the FG stressor with a dopamine D1 agonist (SKF 81297) or antagonist (SCH 23390). Unfortunately, most of the animals were unable to tolerate the drug combination. We lowered the dose of the stressor to 1mg/kg to increase the sample size but found that the low dose was insufficient to alter the animals’ normal range of behavior.

**Figure S3.**
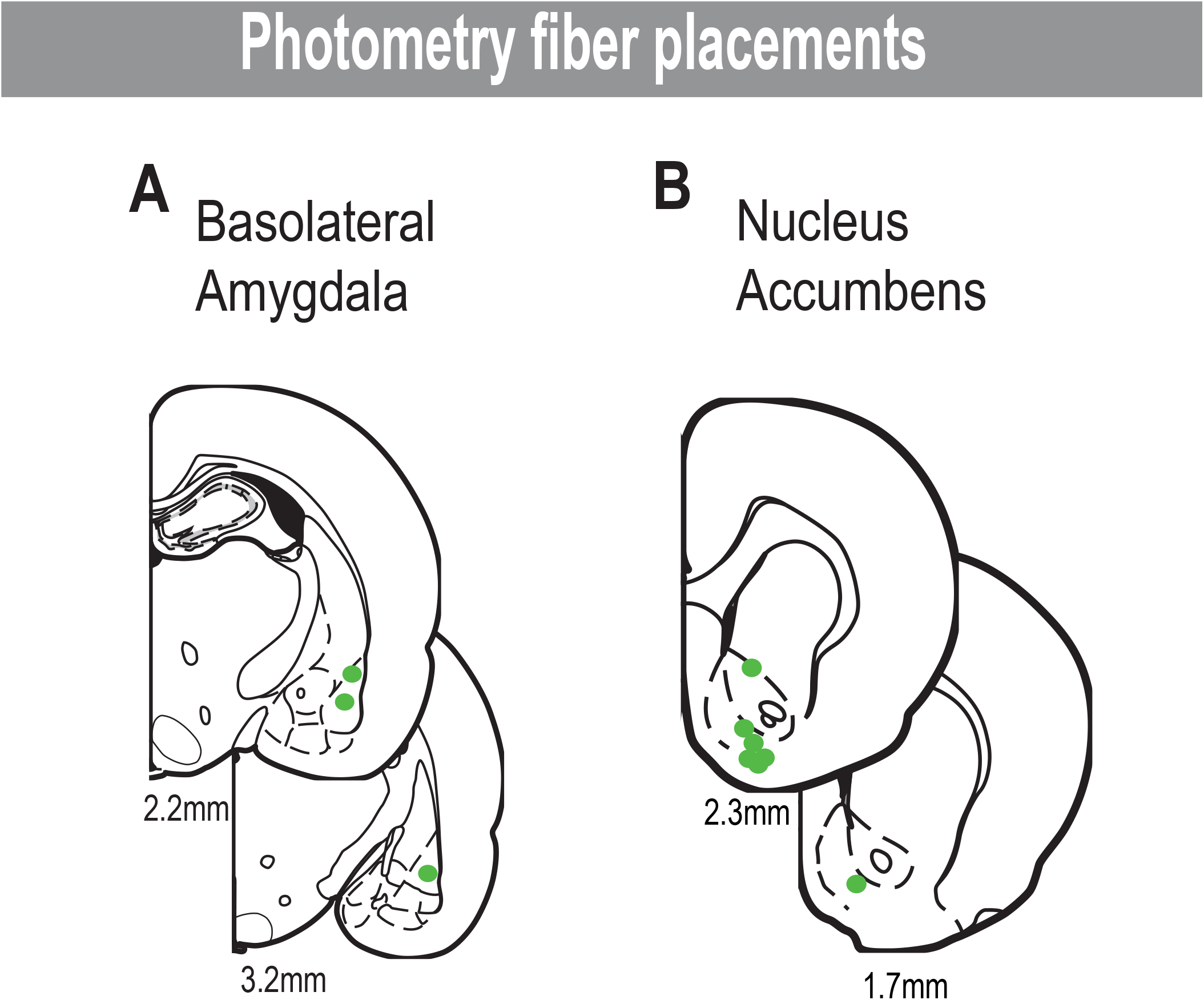
Photometry fiber placement in nucleus basolateral amygdala and nucleus accumbens. A. Green dots in coronal sections represent tips of the fibers used to record noradrenaline dynamics in amygdala. B. Green in coronal sections represent tips of the fibers used to record dopamine dynamics in nucleus accumbens.

**Figure S4.**
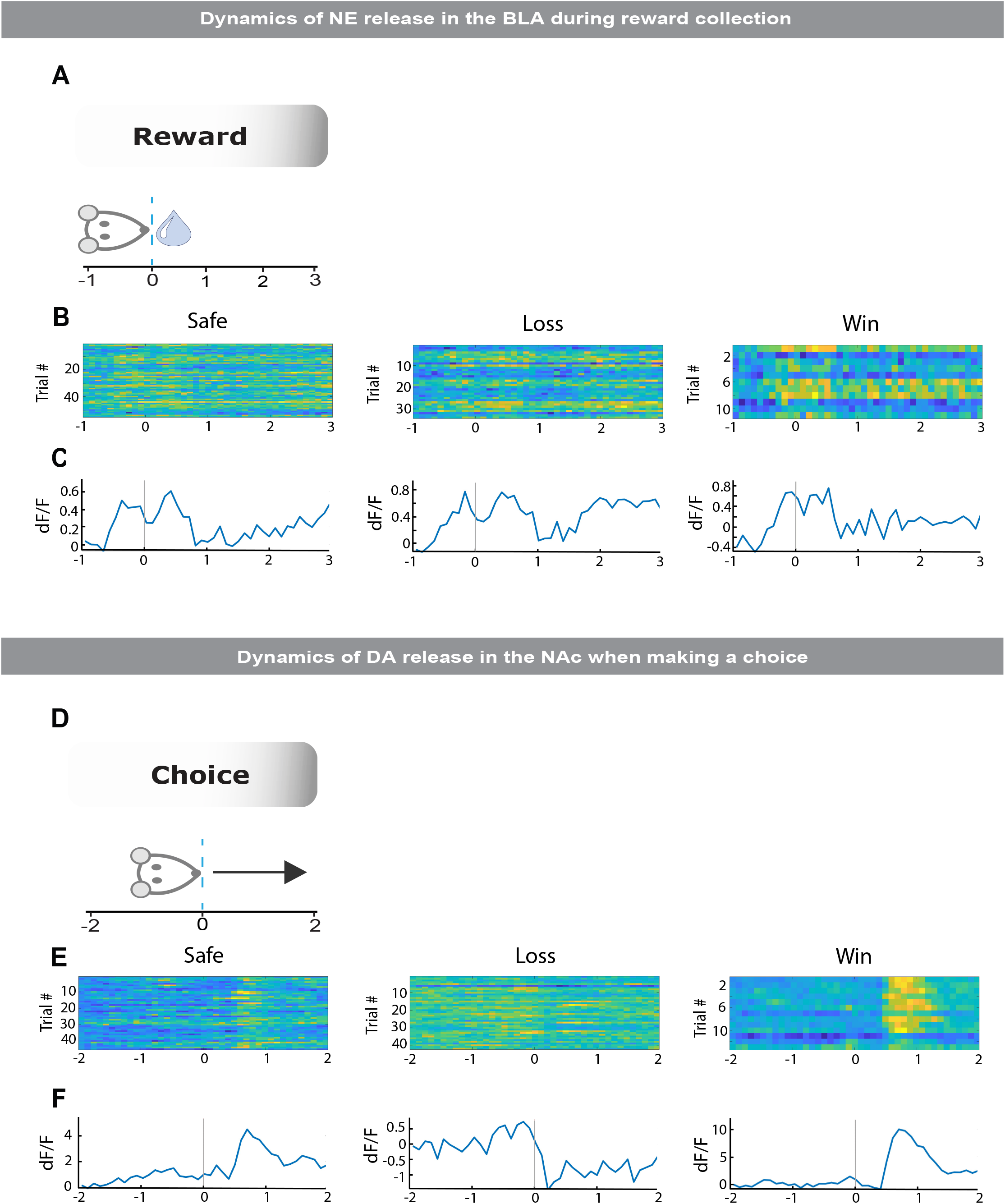
Dynamics of dopamine and noradrenaline signal when aligned to choice response and reward collection in a single animal. A. Schematic representation of reward collection phase. Rat entered the magazine to collect the reward. B. Peri-event time histogram visualize noradrenaline activity in each trial during safe, loss and win collection. C. Averaged change of fluorescence intensity during safe, loss and win collection. D. Schematic representation of choice phase. During choice phase rat approached the screen, touched the screen, and moved towards reward collection. E. Peri-event time histogram visualize dopamine activity in each trial during safe, loss and win choice. F. Averaged change of fluorescence intensity during safe, loss and win choice.

**Figure S5.**
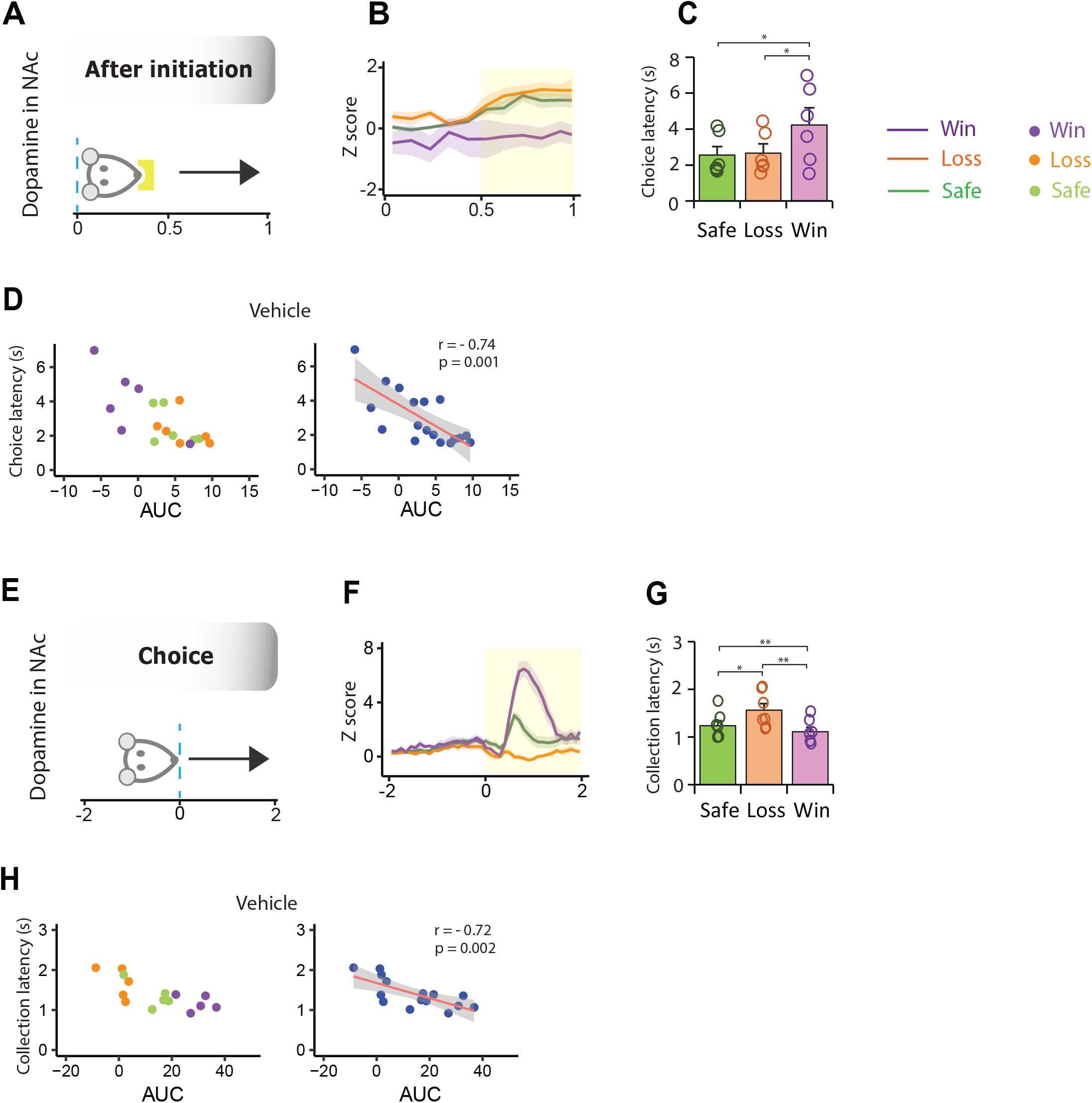

Dopamine activity negatively correlate with choice and collection latencies. A. Schematic representation of initiation phase. After initiation rat moves towards touchscreen for the next choice. B. Mean dopamine signal after initiation (n = 5). Yellow area assigns the period used for signal quantification. C. Choice latency, time between initiation and choice, was affected by the previous trial outcome, as after win rat were slower to make next choice (F1,5= 14.535, p = 0.01; Win vs Safe, p = 0.032; Win vs Loss, p = 0.04). D. Correlation between dopamine activity (area under the curve) after trial initiation and choice latency (r = -0.744, p = 0.001). E. Schematic representation of choice phase. During choice phase rat approached the screen, touched the screen and moved towards reward collection. F. Mean dopamine signal during choice phase (n = 5). Yellow area assigns the period used for signal quantification. G. Collection latency differentiated based on the reward outcome. After a reward win, animals were fast to collect reward, whereas after reward loss, rats took longer to collect the reward (F2,10 = 41.865, p < 0.001; Win vs Safe, p = 0.005; Win vs Loss, p = 0.002; Safe vs Loss, p = 0.012). H. Correlation between dopamine activity (area under the curve, AUC) during choice response and reward collection latency (r = - 0.72, p = 0.002).

## REFERENCES

1. Platt, M.L. and S.A. Huettel, Risky business: the neuroeconomics of decision making under uncertainty. Nat Neurosci, 2008. 11(4): p. 398–403.

2. Croy, M.I. and R.N. Hughes, *feeding behaviour and on the acquisition of learned foraging skills by the fifteen-spined stickleback, Spinachia spinachia L*. Animal Behaviour, 1991. 41(1): p. 161–170.

3. Young, R.J., H. Clayton, and C.J. Barnard, Risk-sensitive foraging in bitterlings, Rhodeus sericus: effects of food requirement and breeding site quality. Animal Behaviour, 1990. 40(2): p. 288–297.

4. Caraco, T., et al., Risk-sensitivity: ambient temperature affects foraging choice. Animal Behaviour, 1990. 39(2): p. 338–345.

5. Case, D.A., P. Nichols, and E. Fantino, Pigeons’ preference for variable-interval water reinforcement under widely varied water budgets. J Exp Anal Behav, 1995. 64(3): p. 299–311.

6. Harder, L.D. and L.A. Real, Why are Bumble Bees Risk. Ecology, 1987. 68(4): p. 1104–1108.

7. Waddington, K.D., Factors Influencing Pollen Flow in Bumblebee-Pollinated Delphinium virescens. Oikos, 1981. 37(2): p. 153–159.

8. Santos, L.R. and A.G. Rosati, The evolutionary roots of human decision making. Annu Rev Psychol, 2015. 66: p. 321–47.

9. Hermans, E.J., et al., Stress-related noradrenergic activity prompts large-scale neural network reconfiguration. Science, 2011. 334(6059): p. 1151–3.

10. Margittai, Z., et al., Exogenous cortisol causes a shift from deliberative to intuitive thinking. Psychoneuroendocrinology, 2016. 64: p. 131–5.

11. Margittai, Z., et al., Combined Effects of Glucocorticoid and Noradrenergic Activity on Loss Aversion. Neuropsychopharmacology, 2018. 43(2): p. 334–341.

12. Jones, C.M. and E.F. McCance-Katz, Co-occurring substance use and mental disorders among adults with opioid use disorder. Drug Alcohol Depend, 2019. 197: p. 78–82.

13. Starcke, K. and M. Brand, Decision making under stress: a selective review. Neurosci Biobehav Rev, 2012. 36(4): p. 1228–48.

14. Wemm, S.E. and R. Sinha, Drug-induced stress responses and addiction risk and relapse. Neurobiol Stress, 2019. 10: p. 100148.

15. Arnsten, A.F., Stress signalling pathways that impair prefrontal cortex structure and function. Nat Rev Neurosci, 2009. 10(6): p. 410–22.

16. Buffalari, D.M. and A.A. Grace, Noradrenergic modulation of basolateral amygdala neuronal activity: opposing influences of alpha-2 and beta receptor activation. J Neurosci, 2007. 27(45): p. 12358–66.

17. Campioni, M.R., M. Xu, and D.S. McGehee, Stress-induced changes in nucleus accumbens glutamate synaptic plasticity. J Neurophysiol, 2009. 101(6): p. 3192–8.

18. McEwen, B.S., C. Nasca, and J.D. Gray, Stress Effects on Neuronal Structure: Hippocampus, Amygdala, and Prefrontal Cortex. Neuropsychopharmacology, 2016. 41(1): p. 3–23.

19. Abela, A.R. and Y. Chudasama, Noradrenergic alpha2A-receptor stimulation in the ventral hippocampus reduces impulsive decision-making. Psychopharmacology (Berl), 2014. 231(3): p. 521–31.

20. Floresco, S.B., et al., Differential Contributions of Nucleus Accumbens Subregions to Cue-Guided Risk/Reward Decision Making and Implementation of Conditional Rules. J Neurosci, 2018. 38(8): p. 1901–1914.

21. Ghods-Sharifi, S., J.R. St Onge, and S.B. Floresco, Fundamental contribution by the basolateral amygdala to different forms of decision making. J Neurosci, 2009. 29(16): p. 5251–9.

22. St Onge, J.R. and S.B. Floresco, Prefrontal cortical contribution to risk-based decision making. Cereb Cortex, 2010. 20(8): p. 1816–28.

23. Tremblay, M., et al., Dissociable effects of basolateral amygdala lesions on decision making biases in rats when loss or gain is emphasized. Cogn Affect Behav Neurosci, 2014. 14(4): p. 1184–95.

24. Galvez, R., M.H. Mesches, and J.L. McGaugh, Norepinephrine release in the amygdala in response to footshock stimulation. Neurobiol Learn Mem, 1996. 66(3): p. 253–7.

25. Hatfield, T., C. Spanis, and J.L. McGaugh, Response of amygdalar norepinephrine to footshock and GABAergic drugs using in vivo microdialysis and HPLC. Brain Res, 1999. 835(2): p. 340–5.

26. Ferry, B., P.J. Magistretti, and E. Pralong, Noradrenaline modulates glutamate-mediated neurotransmission in the rat basolateral amygdala in vitro. Eur J Neurosci, 1997. 9(7): p. 1356–64.

27. Onur, O.A., et al., Noradrenergic enhancement of amygdala responses to fear. Soc Cogn Affect Neurosci, 2009. 4(2): p. 119–26.

28. Xu, P., et al., Amygdala-prefrontal connectivity modulates loss aversion bias in anxious individuals. Neuroimage, 2020. 218: p. 116957.

29. Imperato, A., et al., Stress-induced enhancement of dopamine and acetylcholine release in limbic structures: role of corticosterone. Eur J Pharmacol, 1989. 165(2-3): p. 337–8.

30. Kalivas, P.W. and P. Duffy, Selective activation of dopamine transmission in the shell of the nucleus accumbens by stress. Brain Res, 1995. 675(1-2): p. 325–8.

31. Piazza, P.V., et al., Glucocorticoids have state-dependent stimulant effects on the mesencephalic dopaminergic transmission. Proc Natl Acad Sci U S A, 1996. 93(16): p. 8716–20.

32. Salamone, J.D. and M. Correa, The mysterious motivational functions of mesolimbic dopamine. Neuron, 2012. 76(3): p. 470–85.

33. Salamone, J.D., et al., Effort-related functions of nucleus accumbens dopamine and associated forebrain circuits. Psychopharmacology (Berl), 2007. 191(3): p. 461–82.

34. Wanat, M.J., A. Bonci, and P.E. Phillips, CRF acts in the midbrain to attenuate accumbens dopamine release to rewards but not their predictors. Nat Neurosci, 2013. 16(4): p. 383–5.

35. Shafiei, N., et al., Acute stress induces selective alterations in cost/benefit decision-making. Neuropsychopharmacology, 2012. 37(10): p. 2194–209.

36. Evans, A.K. and C.A. Lowry, Pharmacology of the beta-carboline FG-7,142, a partial inverse agonist at the benzodiazepine allosteric site of the GABA A receptor: neurochemical, neurophysiological, and behavioral effects. CNS Drug Rev, 2007. 13(4): p. 475–501.

37. McGregor, I.S. and D.M. Atrens, Stressor-like effects of FG-7142 on medial prefrontal cortex self-stimulation. Brain Res, 1990. 516(1): p. 170–4.

38. Murphy, B.L., et al., Dopamine and spatial working memory in rats and monkeys: pharmacological reversal of stress-induced impairment. J Neurosci, 1996. 16(23): p. 7768–75.

39. Bruno, C.A., et al., pMAT: An open-source software suite for the analysis of fiber photometry data. Pharmacol Biochem Behav, 2021. 201: p. 173093.

40. D., K. and T. A., Prospect Theory: An Analysis of Decision under Risk. Econometrica, 1979. 47: p. 263–292.

41. Franowicz, J.S. and A.F. Arnsten, The alpha-2a noradrenergic agonist, guanfacine, improves delayed response performance in young adult rhesus monkeys. Psychopharmacology (Berl), 1998. 136(1): p. 8–14.

42. Kim, S., et al., Effects of alpha-2A adrenergic receptor agonist on time and risk preference in primates. Psychopharmacology (Berl), 2012. 219(2): p. 363–75.

43. Clark, C.A. and A. Dagher, The role of dopamine in risk taking: a specific look at Parkinson’s disease and gambling. Front Behav Neurosci, 2014. 8: p. 196.

44. DeVito, E.E., et al., A preliminary study of the neural effects of behavioral therapy for substance use disorders. Drug Alcohol Depend, 2012. 122(3): p. 228–35.

45. Maier, S.F. and L.R. Watkins, Stressor controllability and learned helplessness: the roles of the dorsal raphe nucleus, serotonin, and corticotropin-releasing factor. Neurosci Biobehav Rev, 2005. 29(4-5): p. 829–41.

46. Horger, B.A. and R.H. Roth, The role of mesoprefrontal dopamine neurons in stress. Crit Rev Neurobiol, 1996. 10(3-4): p. 395–418.

47. McCullough, L.D. and J.D. Salamone, Involvement of nucleus accumbens dopamine in the motor activity induced by periodic food presentation: a microdialysis and behavioral study. Brain Res, 1992. 592(1-2): p. 29–36.

48. Tanaka, H., et al., Aging, habitual exercise, and dynamic arterial compliance. Circulation, 2000. 102(11): p. 1270–5.

49. Winstanley, C.A., Gambling rats: insight into impulsive and addictive behavior. Neuropsychopharmacology, 2011. 36(1): p. 359.

50. Asan, E., The catecholaminergic innervation of the rat amygdala. Adv Anat Embryol Cell Biol, 1998. 142: p. 1-118.

51. de Kloet, E.R., M. Joels, and F. Holsboer, Stress and the brain: from adaptation to disease. Nat Rev Neurosci, 2005. 6(6): p. 463–75.

52. Herman, J.P., et al., Limbic system mechanisms of stress regulation: hypothalamo-pituitary-adrenocortical axis. Prog Neuropsychopharmacol Biol Psychiatry, 2005. 29(8): p. 1201–13.

53. Piazza, P.V. and M. Le Moal, Glucocorticoids as a biological substrate of reward: physiological and pathophysiological implications. Brain Res Brain Res Rev, 1997. 25(3): p. 359–72.

54. Eshel, N., et al., Striatal dopamine integrates cost, benefit, and motivation. Neuron, 2024. 112(3): p. 500–514 e5.

55. Salamone, J.D., et al., Beyond the reward hypothesis: alternative functions of nucleus accumbens dopamine. Curr Opin Pharmacol, 2005. 5(1): p. 34–41.

56. Arnsten, A.F. and F.M. Leslie, Behavioral and receptor binding analysis of the alpha 2-adrenergic agonist, 5-bromo-6 [2-imidazoline-2-yl amino] quinoxaline (UK-14304): evidence for cognitive enhancement at an alpha 2-adrenoceptor subtype. Neuropharmacology, 1991. 30(12A): p. 1279-89.

57. Arnsten, A.F., et al., Noradrenergic influences on prefrontal cortical cognitive function: opposing actions at postjunctional alpha 1 versus alpha 2-adrenergic receptors. Adv Pharmacol, 1998. 42: p. 764–7.

58. Connolly, N.P., et al., A test of stress, cues, and re-exposure to large wins as potential reinstaters of suboptimal decision making in rats. Front Psychol, 2015. 6: p. 394.

59. Sun, H., et al., Yohimbine increases impulsivity through activation of cAMP response element binding in the orbitofrontal cortex. Biol Psychiatry, 2010. 67(7): p. 649–56.

60. Tabbara, R.I., et al., The pharmacological stressor yohimbine, but not U50,488, increases responding for conditioned reinforcers paired with ethanol or sucrose. Psychopharmacology (Berl), 2020. 237(12): p. 3689–3702.

61. Stanford, S.C., Central noradrenergic neurones and stress. Pharmacol Ther, 1995. 68(2): p. 297–42.

62. LeDoux, J.E., Emotion circuits in the brain. Annu Rev Neurosci, 2000. 23: p. 155–84.

63. McGaugh, J.L., The amygdala modulates the consolidation of memories of emotionally arousing experiences. Annu Rev Neurosci, 2004. 27: p. 1–28.

64. Mantz, J., et al., Differential effects of ascending neurons containing dopamine and noradrenaline in the control of spontaneous activity and of evoked responses in the rat prefrontal cortex. Neuroscience, 1988. 27(2): p. 517–26.

65. T., N., et al., Alpha-2 adrenoceptor mediated inhibition of [3H] noradrenaline release from rat hippocampal synaptosomes: potentiation by chronic treatment with desipramine or electroconvulsive shock. European Journal of Pharmacology, 1983. 86(3-4): p. 349–356.

66. Tsuda, A., et al., [Stressor controllability and brain noradrenaline turnover in rats]. Yakubutsu Seishin Kodo, 1987. 7(3): p. 363–74.

67. Bremner, J.D., et al., Decreased benzodiazepine receptor binding in prefrontal cortex in combat-related posttraumatic stress disorder. Am J Psychiatry, 2000. 157(7): p. 1120–6.

68. Weizman, R., et al., Repeated swim stress alters brain benzodiazepine receptors measured in vivo. J Pharmacol Exp Ther, 1989. 249(3): p. 701–7.

69. Baba, H., et al., Norepinephrine facilitates inhibitory transmission in substantia gelatinosa of adult rat spinal cord (part 2): effects on somatodendritic sites of GABAergic neurons. Anesthesiology, 2000. 92(2): p. 485–92.

70. Li, R., et al., Synapses on GABAergic neurons in the basolateral nucleus of the rat amygdala: double-labeling immunoelectron microscopy. Synapse, 2002. 43(1): p. 42–50.

71. St Onge, J.R. and S.B. Floresco, Dopaminergic modulation of risk-based decision making. Neuropsychopharmacology, 2009. 34(3): p. 681–97.

72. van Gaalen, M.M., et al., Behavioral disinhibition requires dopamine receptor activation. Psychopharmacology (Berl), 2006. 187(1): p. 73–85.

73. Ljungberg, T. and M. Enquist, Effects of dopamine D-1 and D-2 antagonists on decision making by rats: no reversal of neuroleptic-induced attenuation by scopolamine. J Neural Transm Gen Sect, 1990. 82(3): p. 167–79.

74. Simon, N.W., et al., Dopaminergic modulation of risky decision-making. J Neurosci, 2011. 31(48): p. 17460–70.

75. Zeeb, F.D. and C.A. Winstanley, Functional disconnection of the orbitofrontal cortex and basolateral amygdala impairs acquisition of a rat gambling task and disrupts animals’ ability to alter decision-making behavior after reinforcer devaluation. J Neurosci, 2013. 33(15): p. 6434–43.

